# A metabolic life table exemplified by sockeye salmon

**DOI:** 10.1101/2020.03.13.990986

**Authors:** Joseph R. Burger, Chen Hou, Charles A.S Hall, James H. Brown

## Abstract

1. A metabolic life table (MLT) is a combination of energy budget and life table that quantifies metabolism and life history over an entire life cycle. It provides a conceptual framework for integrating data on physiology, demography and ecology that are usually the subject of discipline- and taxon-specific studies.
2. Our MLT for sockeye salmon revives John Brett’s classic data on metabolism, growth, survival and reproduction to provide a synthetic analysis of metabolic performance and its life history consequences.
3. The MLT quantifies the energy budget of an average female over her life cycle. The early stages in fresh water have low rates of growth and mortality. Then juveniles enter the ocean, feed voraciously, grow rapidly, and accumulate a store of biomass; More than 98% of the total lifetime assimilation and growth occurs in the ocean. Maturing adults stop feeding, return to fresh water, expend stored body energy to fuel migration and reproduction, and leave a clutch of eggs and a depleted carcass.
4. The MLT also quantifies the energetic contribution of salmon to freshwater and marine ecosystems. Salmon are very efficient: of the food energy in their zooplankton prey, 47% is expended on respiration to fuel activity, 23% is allocated to growth and reproduction to produce biomass, and 30% is excreted in feces. Although 96% of lifetime biomass production occurs in the ocean, about 29% is transported into fresh water in the bodies of maturing adults, where their carcasses and gametes provide an important “marine subsidy” to the energy and nutrient budgets of freshwater and terrestrial ecosystems.
5. The MLT highlights some features of the salmon metabolism – variation in rates of assimilation, respiration and production with body size and temperature – that are qualitatively similar to other ectotherms and predictions of metabolic theory. But because of the unusual physiology, life history, ecosystem impacts and socio-economic importance of wild-caught salmon, reanalyzing Brett’s data in the context of a MLT has additional broad applications for basic and applied ecology.

## Introduction

A challenge of contemporary basic and applied ecology is to synthesize the detailed information available in specialized databases and journals to elucidate general patterns and processes. Empirical and theoretical metabolic ecology use energy as a common currency to make explicit linkages among life history, ecology and evolution (Brown, Gillooly, Allen, Savage, & West, 2004; Burger, Hou, & Brown, 2019; Hall, 1972; Kooijman & Kooijman, 2010; Lotka, 1922; Odum & Pinkerton, 1955; Sibly, Brown, & Kodric-Brown, 2012; Van Valen, 1976). These connect anatomy, physiology and behavior at the level of individual organisms to ecological patterns and processes at population, community and ecosystem levels. They also have important applications to conservation and management. Many studies apply short-term measurements of resting, active and field metabolism of particular species to address questions about the ecological costs and benefits and the evolutionary consequences of individual organism-level traits and ecosystem-level phenomena (e.g., Peters 1983; Brown et al. 2004; Sibly et al. 2012). Largely missing, however, are comprehensive studies of single species that can address of how metabolic energy is acquired and expended over an entire life cycle and interpret them in ecological and evolutionary contexts. Pacific salmon (*Oncorhynchus spp.*) provide an excellent model organism for two reasons: their semelparous life history simplifies quantification, obviating the need for detailed data and analyses to address ideterminate growth and semelparous reproduction (Fig. 1); and they have been studied intensively – long prized as food for humans, wild salmon fisheries are worth billions of dollars – so there are relevant physiological and demographic data for the entire life cycle (Knapp, Guettabi, & Goldsmith, 2013; Quinn, 2018).

**Figure 1.**
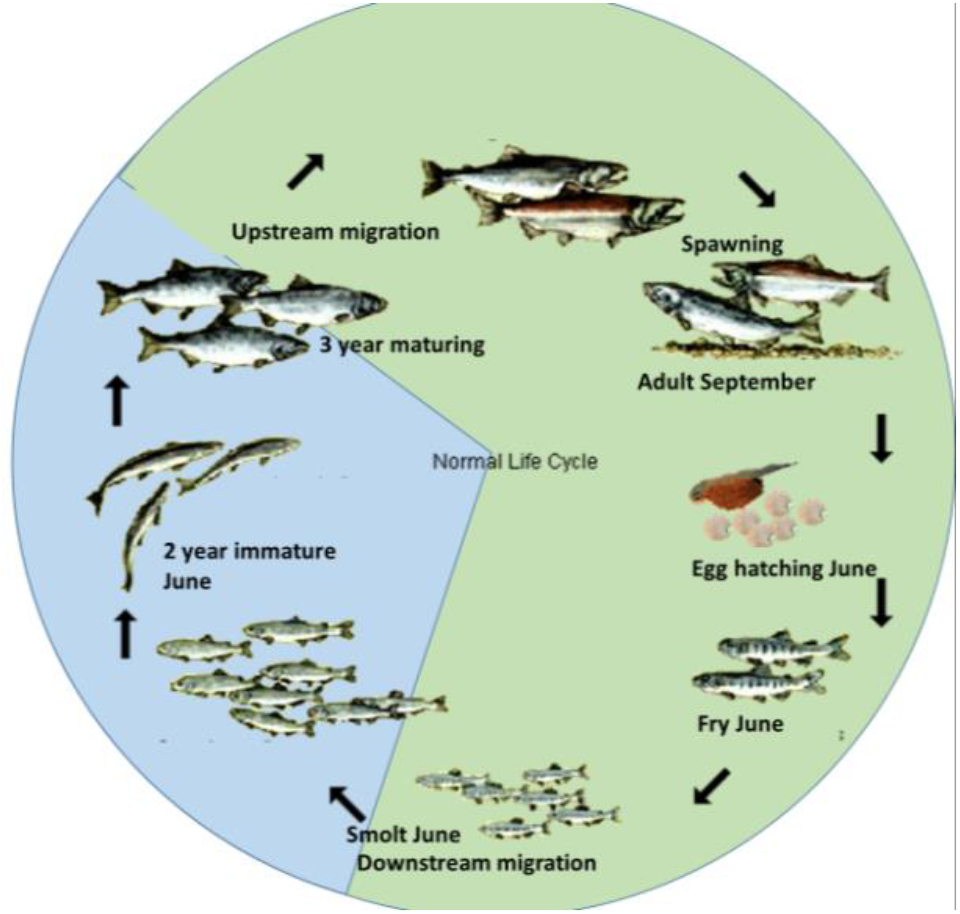
Life cycle of sockeye salmon after Brett (JOHN R. Brett, 1983). The Babine Lake population is semelparous – expending energy on a single, end-of-life reproductive effort, and anadromous – spending about equal time – two years – reproducing in fresh water (green) and feeding and growing in the ocean (blue). Modified from (“Https://Www.Google.Com/Search?Q=salmon+life+cycle&rlz=1C1CHFX_enUS555US555&source=lnms&tbm=isch&sa=X&ved=0ahUKEwj7rZiC3PXWAhWBLmMKHRIiAM8Q_AUICigB&biw=1067&bih=423#imgrc=WiztTrFdqAKvPM,” n.d.) and not drawn to scale.

Canadian fisheries biologist John “Roly” Brett introduced a “life table of energetics” to synthesize his life’s work on the physiological ecology of the sockeye salmon (*Oncorhynchus nerka*) that breed in Babine Lake, British Columbia (Brett, 1986; Brett, 1983). His goal was to “put together a comprehensive estimate of the life energetics of an average sockeye salmon that commences life as a fertilized egg weighing 0.13 g containing 372 cal … and terminates growth as a 2,270 g adult containing 4,200 kcal when entering fresh water on its final journey” (Brett, 1983), p.30). Brett adopted the form of a standard life table to present his data on body size, energy content, growth and metabolism as a function of age over an entire life cycle (Table 1). Here we resurrect and extend Brett’s concept of a “life table of energetics” to construct a metabolic life table (MLT) that combines the demography of a traditional life table with the mass-energy balance of physiological ecology. By quantifying energy allocation to components of fitness – survival, growth and reproduction – as a function of age, the MLT highlights the dual roles of universal constraints and special adaptations in metabolic ecology and life history of salmon. We demonstrate the practical utility and broad applicability of the MLT approach, and show how it goes beyond traditional demography and short-term studies of metabolism to provide a synthetic picture of the central role of energy in physiology, behavior, ecology and evolution.

**Table 1.** Metabolic life table (MLT) for sockeye salmon in Babine Lake, based on data in Brett (1983, 1986). The definitions of stages and traits follow Brett, although we have transformed his data into modern units and calculated additional parameters. Green rows represent stages in freshwater and blue rows marine. Unlike some life tables that are based only on females, this one is based on replacement of a male and female parent at steady state.

| (°C) | Stage<br>(Months) | Wet<br>weight<br>(g) | Numbe<br>r alive<br>number | Body<br>energy<br>content<br>(kJ) | Productio<br>n rate<br>(watts) | Whole-organism<br>respiration<br>rate<br>(watts) | Assimilati<br>on rate<br>(watts) | Energy<br>expende<br>d<br>(kJ) | Cumulativ<br>e energy<br>expended<br>(kJ) | Standing<br>stock of<br>biomass<br>(kJ) | Cohort<br>production<br>lost to<br>mortality<br>(kJ) | Cohort<br>respiration<br>before dying<br>(kJ) | Cohort<br>assimilation<br>before dying<br>(kJ) |
| --- | --- | --- | --- | --- | --- | --- | --- | --- | --- | --- | --- | --- | --- |
| 6 | Egg (0) | 0.013 | 3000 | 0.13 | 0.0000 | 0.0000002 | 0.0000000 | 0.00 | 0.00 | 386.03 | 0.00 | 0.00 | 0.00 |
| 6.0 | Fry (9) | 0.25 | 600 | 1.15 | 0.00 | 0.0002144 | 0.0009733 | 0.08 | 2.09 | 690.69 | 1535.79 | 2511.6 | 4047.39 |
| 11.0 | Fry (11) | 1.00 | 367 | 4.97 | 0.00 | 0.00078 | 0.0021052 | 2.09 | 9.21 | 1825.15 | 713.42 | 1316.623794 | 2030.04 |
| 9.0 | Fry (13) | 3.20 | 224 | 16.73 | 0.00 | 0.00233 | 0.0060826 | 7.12 | 34.74 | 3755.26 | 1546.30 | 3131.982487 | 4678.29 |
| 4.0 | Fry (17) | 4.50 | 137 | 23.51 | 0.00 | 0.00246 | 0.0032261 | 25.53 | 54.00 | 3228.30 | 1753.78 | 3868.102532 | 5621.89 |
| 5.0 | Smolt yearling<br>(20) | 5.00 | 84 | 26.00 | 0.00 | 0.00254 | 0.0029052 | 19.26 | 62.37 | 2183.59 | 1319.87 | 3102.698087 | 4422.57 |
| 11.0 | Smolt yearling<br>(21) | 5.40 | 75 | 29.39 | 0.00 | 0.00366 | 0.0045643 | 8.37 | 84.56 | 2195.64 | 257.02 | 681.9018036 | 938.93 |
| 12.0 | Smolt yearling<br>(22) | 15.00 | 66 | 86.34 | 0.03 | 0.02006 | 0.0547624 | 22.19 | 179.16 | 5738.04 | 477.72 | 1088.681233 | 1566.40 |
| 12.5 | Smolt yearling<br>(23) | 50.00 | 59 | 304.74 | 0.14 | 0.06203 | 0.2060148 | 94.60 | 400.60 | 18015.50 | 1436.05 | 2128.900796 | 3564.95 |
| 11.0 | Smolt yearling<br>(24) | 110.00 | 53 | 708.65 | 0.22 | 0.10927 | 0.3250237 | 221.44 | 733.39 | 37264.19 | 3310.00 | 3703.907398 | 7013.91 |
| 9.0 | Smolt yearling<br>(25) | 200.00 | 47 | 1288.45 | 0.30 | 0.15216 | 0.4489740 | 332.79 | 1117.24 | 60266.29 | 5802.26 | 5376.71596 | 11178.97 |
| 6.5 | Smolt yearling<br>(26) | 250.00 | 42 | 1670.21 | 0.14 | 0.13447 | 0.2775480 | 383.86 | 1438.31 | 69490.30 | 7646.08 | 6604.313975 | 14250.39 |
| 4.8 | Smolt yearling<br>(27) | 290.00 | 37 | 1937.45 | 0.11 | 0.11523 | 0.2251318 | 321.07 | 1740.54 | 71701.40 | 8293.05 | 7307.319567 | 15600.37 |
| 4.5 | Smolt yearling<br>(28) | 300.00 | 33 | 2004.26 | 0.03 | 0.10903 | 0.1345524 | 302.23 | 2037.33 | 65977.58 | 8059.68 | 7724.674396 | 15784.36 |
| 4.5 | Smolt yearling<br>(29) | 310.00 | 29 | 2071.07 | 0.03 | 0.11266 | 0.1390374 | 296.79 | 2314.44 | 60643.22 | 7412.10 | 7914.886782 | 15326.98 |
| 4.5 | Smolt yearling<br>(30) | 320.00 | 26 | 2137.87 | 0.03 | 0.11630 | 0.1435225 | 277.11 | 2666.48 | 55682.15 | 6809.22 | 8058.131768 | 14867.35 |
| 5.3 | Smolt yearling (31) | 350.00 | 23 | 2433.74 | 0.09 | 0.14925 | 0.2352539 | 352.04 | 3134.06 | 56383.71 | 6578.69 | 8347.156985 | 14925.85 |
| 7.3 | Smolt yearling (32) | 380.00 | 21 | 2670.67 | 0.08 | 0.21913 | 0.3026008 | 467.58 | 3813.86 | 55035.73 | 6533.73 | 8893.456575 | 15427.18 |
| 9.0 | 2 yr immature (33) | 420.00 | 18 | 2962.01 | 0.11 | 0.29511 | 0.4082625 | 679.81 | 4715.95 | 54294.69 | 6413.22 | 9711.818069 | 16125.04 |
| 11.0 | 2 yr immature (34) | 480.00 | 16 | 3374.75 | 0.18 | 0.41170 | 0.5874999 | 902.08 | 5986.40 | 55024.72 | 6417.62 | 10838.90796 | 17256.53 |
| 12.0 | 2 yr immature (35) | 590.00 | 15 | 4151.67 | 0.33 | 0.54893 | 0.8805507 | 1270.45 | 6142.54 | 60212.21 | 6780.18 | 10926.33887 | 17706.51 |
| 11.5 | 2 yr immature (36) | 720.00 | 13 | 5074.27 | 0.39 | 0.62453 | 1.0122159 | 1562.63 | 7731.96 | 65460.64 | 7392.79 | 11117.6938 | 18510.48 |
| 9.3 | 2 yr immature (37) | 890.00 | 11 | 6263.93 | 0.51 | 0.60379 | 1.1186258 | 1589.42 | 9151.43 | 71878.52 | 8081.41 | 12033.79686 | 20115.20 |
| 6.5 | 2 yr immature (38) | 1000.00 | 10 | 7283.64 | 0.32 | 0.46520 | 0.7883632 | 1419.47 | 10315.56 | 74344.03 | 8589.14 | 12342.04889 | 20931.19 |
| 5.0 | 2 yr immature (39) | 1060.00 | 9 | 7587.54 | 0.17 | 0.43147 | 0.5983565 | 1164.13 | 11380.48 | 68888.12 | 8386.47 | 12235.29054 | 20621.76 |
| 4.5 | 2 yr immature (40) | 1100.00 | 8 | 7873.87 | 0.11 | 0.36247 | 0.4718452 | 1064.92 | 12349.96 | 63588.21 | 7755.83 | 11903.78294 | 19659.62 |
| 4.5 | 2 yr immature (41) | 1130.00 | 7 | 8088.61 | 0.09 | 0.36688 | 0.4526258 | 969.48 | 13257.06 | 58104.24 | 7122.38 | 11425.72549 | 18548.10 |
| 4.5 | 2 yr immature (42) | 1180.00 | 6 | 8594.70 | 0.14 | 0.38311 | 0.5224005 | 907.11 | 14376.40 | 54917.40 | 6621.44 | 10967.44636 | 17588.88 |
| 5.0 | 2 yr immature (43) | 1250.00 | 6 | 9104.55 | 0.20 | 0.44824 | 0.6484892 | 1119.34 | 15805.08 | 51746.79 | 6248.42 | 10655.06793 | 16903.49 |
| 7.0 | 3 yr maturing (44) | 1400.00 | 5 | 10537.00 | 0.46 | 0.67841 | 1.1419302 | 1428.68 | 18086.03 | 53270.57 | 6167.89 | 10642.58165 | 16810.48 |
| 9.5 | 4 yr maturing (45) | 1670.00 | 4 | 12576.42 | 0.86 | 1.07630 | 1.9352652 | 2281.79 | 21685.61 | 56555.24 | 6456.11 | 11109.1341 | 17565.24 |
| 12.5 | Adult-mature (46) | 2100.00 | 4 | 16315.35 | 1.44 | 1.80118 | 3.2365951 | 3598.70 | 26813.00 | 65261.62 | 7178.38 | 12049.84547 | 19228.22 |
| 13.0 | Adult-mature (46.5) | 2270.00 | 4 | 17626.41 | 0.38 | 2.02399 | 2.4081649 | 5127.43 | 29435.53 | 70505.64 | 8485.44 | 14062.13445 | 22547.58 |
| 12.0 | Adult-mature (48) | 2200.00 | 2 | 4455.00 | 0.00 | 2.55858 | 2.5585824 | 2622.53 | 39013.52 | 8910.00 | 6682.50 | 58520.28 | 65202.78 |
| 12.0 | Carcass (48.1) | 2161.00 | 0 | 4169.00 | 0.00 | 0.00000 | 0.0000000 | 5345.52 | 39013.52 | 0.00 | 8338.00 | 78027.04 | 86365.04 |

## Methods

Brett’s “Life Table of Energetics” (Brett, 1983), which gives data on body mass and composition, and energy acquisition and expenditure for 33 stages of a typical four-year life cycle is distilled and modified in Table 1. Brett made some simplifications and assumptions:

1. He assumed a steady state population with reproduction balancing mortality. It is well known that there is considerable variation in the size and composition of the Skeena River runs among years (e.g., Larkin and McDonald 1968). However, in the 1970s and early 1980s the Babine Lake sockeye population and fishery was stable, so we followed Brett (1983, 1986) in assuming steady state.
2. He assumed a simplified four-year life cycle, ignoring the small proportion of the population returning to breed after three or five years.
3. He estimated average values for the population as a whole, ignoring considerable variation among individuals and across years.
4. He used laboratory measurements, simplified models, and corrections for environmental temperature to estimate energy assimilation from feeding and energy expenditure for maintenance and activity.
5. He used additional information from models and other salmon populations to estimate parameters (e.g., growth and mortality) for the marine stages for which direct measurements for the Babine Lake population were not available.
6. He recognized that many parameters were estimated imprecisely, so he rounded off and reported average values.

We added mortality data from (Brett, 1986) to estimate the age structure for the 33 stages in Fig. 2. Using these values — and ignoring seasonal and interannual variation due to movement at sea, seasonal fluctuations in ocean temperature and productivity, and differences among runs — we compiled a MLT for the Babine Lake sockeye population (Table 1). Given the above assumptions, simplifications, and conversions, our MLT is a faithful distillation and synthesis of Brett’s data.

**Figure 2.**
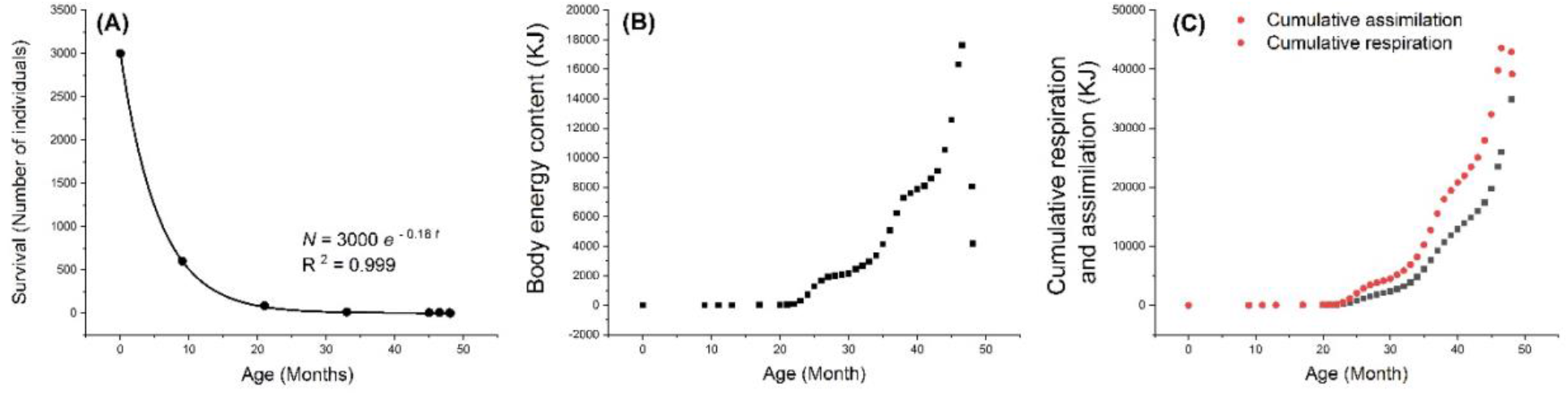
Dynamics of Babine Lake salmon life history and energetics: variation as a function of age. (A) Survival: number alive out of initial clutch of 3,000 eggs. Deviations from the fitted exponential relationship are too small to show on this graph. (B) Body energy content. (C) Cumulative respiration and assimilation.

### Metabolic ecology of sockeye salmon

#### Natural history of the life cycle

Salmon are: 1) anadromous – breeding in fresh water and growing in the ocean, and 2) semelparous – making a single, end-of-life reproductive effort. The life cycle (Fig. 1) starts in September when the mature adults have returned to spawn in tributary streams of Babine Lake. The average female lays 3,000 eggs, each weighing about 0.1 g. Males fertilize the eggs as females deposit them in the streambed. The eggs are buried in gravel and develop over the winter. In June ~600 survivors hatch, move to the lake and feed on plankton. After one year about 84 surviving smolts, each weighing ~5.4 g, migrate downstream and enter the ocean. During the > two years in marine phase, juveniles move between water masses of varying temperature and productivity, feed voraciously on zooplankton, grow rapidly, and accumulate stores of energy-rich lipids and proteins in their distinctive red flesh. In the summer of their fourth year, the maturing adults weigh ~2700 g when they return to coastal waters, stop feeding, and start their upstream migration. By September, they reach the spawning grounds, where females compete to excavate nests in the streambed and males compete to mate with spawning females. Both sexes die after spawning, leaving a clutch of eggs and depleted carcasses, having expended most of their body energy for migration and mating.

#### Life table: survival and reproduction

The first few columns of Table 1 are a life table: a schedule of survival and reproduction as a function of age, stage and body mass. Each generation starts with 3,000 eggs, lasts four years, and ends with another 3,000 eggs spawned by two depleted breeders. In between the eggs hatch, the offspring grow and die as they age, and an average of two survive to breed and continue the life cycle. The dynamics of the life history as a function of age are shown in Fig. 2. Until the salmon stop feeding and enter fresh water for the return migration, cumulative assimilation, respiration and biomass increase with age; deviations from smooth monotonic curves reflect variations in energy income (assimilation from feeding), expenditure (respiration for swimming), and associated variation in temperature (Brett, 1972; Brett & Glass, 1973; Brett, 1983). During the last three months of life, the returning adults use most of their stored body energy on locomotor and reproductive activity, leaving depleted carcasses and a new clutch of eggs.

#### Metabolism: body mass and temperature dependence of respiration, growth and assimilation

The MLT (Table 1) documents variation in rates of respiration, growth and assimilation with age, stage, body mass and environmental temperature at monthly intervals over one generation. We compare these data to predictions from general metabolic theory (Brown et al., 2004; Calder, 1984; Peters, 1986; Schmidt-Nielsen, 1984; Sibly et al., 2012). Whole-organism rates, *B*, characteristically increase with body mass, *m*, scaling as power laws of the form

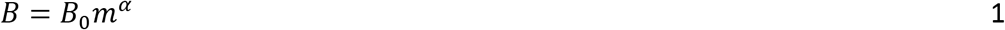

where *B*_o_ is a normalization constant, and the exponent *α* is characteristically close to 3/4 for respiration rate and most other whole-organism rates (Brown et al., 2004; Calder, 1984; Kleiber, 1932; Peters, 1986; Schmidt-Nielsen, 1984; Sibly et al., 2012; West, Brown, & Enquist, 1997). In ectothermic animals, such as salmon, these rates also vary with body temperature with Q_10_ ≈ 2.5 (increasing ~2.5 times for every 10°C increase in temperature (Gillooly, 2001; Huey & Kingsolver, 2019).

Brett’s data for sockeye salmon are qualitatively consistent with predictions from metabolic scaling theory, but with quantitative caveats. Brett (1983, p. 33) determined that the temperature dependence was “equivalent to a Q_l0_ of about 2.3.” We used this value to temperature-correct the rates of respiration, growth and assimilation, and analyze their scalings with body mass (Fig. 3). The scaling relationships are positive as has been found universally, but the exponents (slopes) are consistently higher than α ≈ 0.75 expected from general standard metabolic theory (Brown et al., 2004) and data for most other animals (e.g., “Kleiber’s rule”: (Calder, 1984; Kleiber, 1932; Peters, 1986; Schmidt-Nielsen, 1984; Sibly et al., 2012). We are uncertain how to interpret these deviations. On the one hand, salmon have an unusual physiology and ecology as indicated above. On the other hand, the data might be inaccurate. Brett (Brett, 1983), p. 32) cautions “More often than not when compiling energy budgets growth is measured, metabolism estimated, assimilation efficiency assumed, and excretion deduced…. The components of the balanced equation are sensitive to, and respond differently to, the chief variables of temperature, body size, and activity.” Since we have no objective basis to question or correct Brett’s values, we use them here with this caveat.

**Figure 3.**
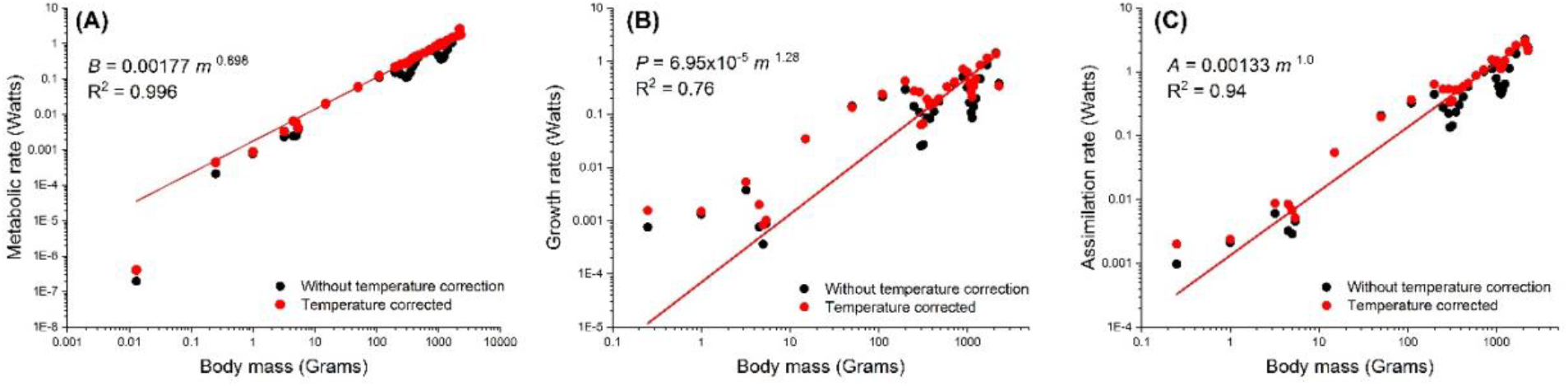
Plots on logarithmic axes showing power-law scalings of rates of whole-organism metabolism as functions of body mass before (black symbols) and after temperature (red symbols and regression lines) correction). (A) Respiration rate (Brett’s “total metabolic rate”): *B* = 0.00177*m*^0.898^; R^2^ = 0.996; B) Rate of biomass production (from Brett’s “growth rate” in %/day *P* = 6.95×10^5^*m*^1.29^; R^2^ = 0.76; (C) Assimilation rate (sum of the above two rates): *A* = 0.00133*m*^1.0^; R^2^ = 0.94.

#### Energy balance of an individual

An energy budget for the average individual surviving to breed accounts for stocks and flows of energy over one generation (Fig. 4):

1. egg: The energy budget starts with a single fertilized egg, weighing 0.10 g and containing 1.25 kJ of energy. The egg lies inactive in the gravel, transforms from an embryo into a fry, and hatches after about nine months, having lost about one-third of its mass and energy in respiration.
2. feeding, maintenance and growth: In the three plus years between hatching and breeding, the offspring assimilates 47,124 kJ of energy by feeding, expends 29,498 kJ (63%) on aerobic respiration, and produces 17,626 kJ (37%) in body energy reserves. More than 98% of the total lifetime assimilation and growth occurs in the ocean, where the juveniles swim and feed almost continuously.
3. migration and breeding: Once a mature salmon enters fresh water, it stops feeding and lives off body reserves. In the last 45 days of life, an individual expends 9,583 kJ (54%) of its body energy on respiration for expensive migration and spawning.
4. parental investment: A fraction of stored body energy is allocated to parental investment in gametes. The clutch of 3,000 eggs contains 3,880 kJ (22% and 8.3% of lifetime biomass production and assimilation, respectively). The male invests less energy, 938 kJ, in sperm.
5. carcass: When the spawned-out adult dies, its weight has decreased from 2,270 to 1,810 g. The carcass, depleted of nearly all fat and much of its protein, contains mostly water. Of the 17,626 kJ stored in the body at the start of migration, 13,463 kJ has been expended, 9,583 kJ on respiration, 4,163 kJ in the depleted carcass and 3,880 kJ on eggs.
6. production efficiency: The efficiency of individual production, *T*_*ind*_, is the ratio of output over input or production over assimilation:

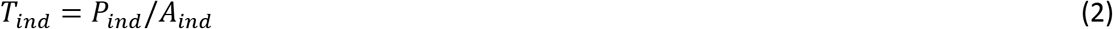

**Figure 4.**
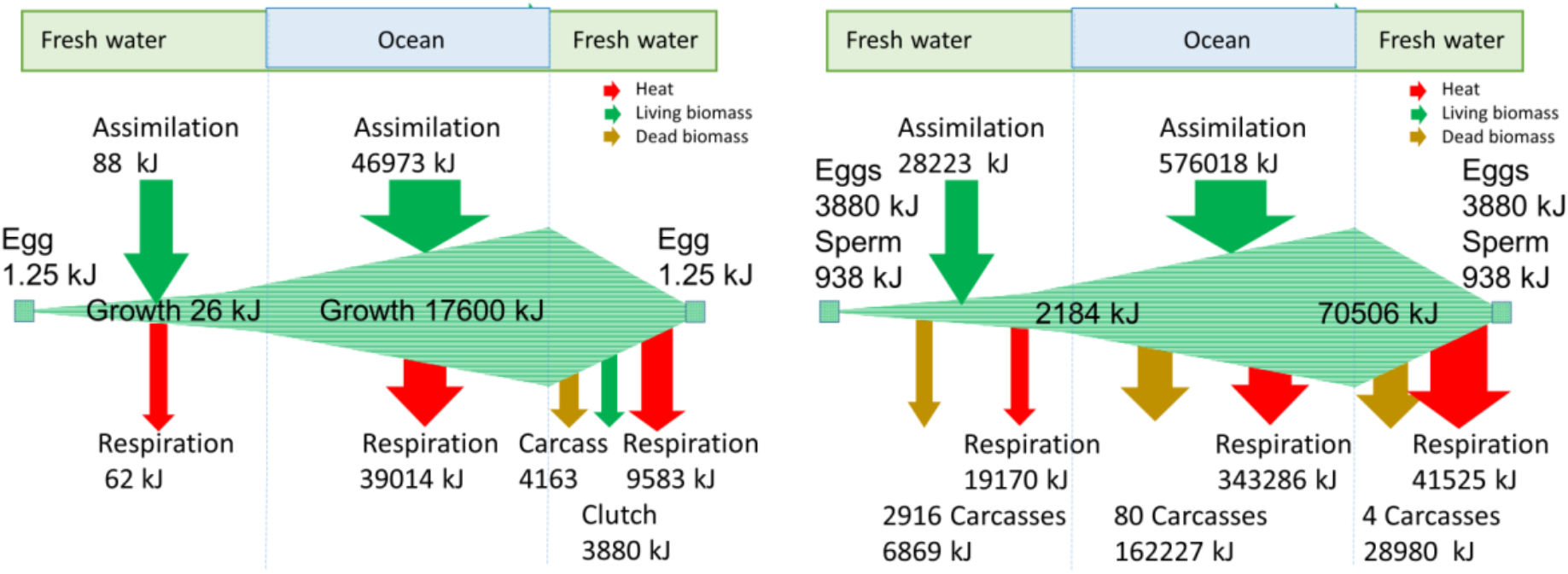
Balanced energy budgets for Babine Lake sockeye salmon: (A) for an individual female. The 4-year life cycle starts and ends with a single egg laid in fresh water; the offspring hatches, grows and enters the ocean, where it grows more rapidly before re-entering fresh water for the return migration; the 17626 kJ of stored body energy is all dissipated in fresh water as respiration, depleted carcass and clutch of eggs. The budget for a male is similar but he allocates less energy to sperm and more to respiration for courtship and mating. (B) for the cohort of all offspring produced by a surviving-to-spawning female. The 3,000 fry suffer high mortality and grow only modesty before entering the ocean. After voracious feeding, rapid growth, and additional mortality at sea, four survivors rapidly deplete their stored body energy on respiration to fuel migration and breeding, leaving a depleted carcass and a new clutch of 3,000 eggs.

It varies with water temperature and age (Table 1), decreasing from >60% for fry and smolts in fresh water to <25% for the older stages in ocean water. By the time a female stops feeding and enters fresh water for the return migration, she has accumulated 37% of assimilated energy in her body as growth. When she dies three months later, after expending more than half of this store on respiration for migration and spawning, her clutch of eggs and depleted body contain just 18% of the energy she assimilated over her lifetime.

#### Energy balance of a cohort

At least as relevant for life history and ecology is the energy budget for the population of a representative cohort (Fig. 4B). This accounting includes intake and expenditure of all offspring produced by a pair of breeders, including those that die before reproducing. Brett (Brett, 1986) called these losses the “life-cycle deficit” to estimate the “food conversion efficiency” of the Babine Lake population.

Since the population was approximately constant during Brett’s study, we assumed steady state and used the MLT (Table 1) to compile a balanced energetic budget for a cohort (Fig. 4B, Table 2):

1. juveniles in fresh water: The energy budget for the cohort starts with 3,000 fertilized eggs containing 3,880 kJ of energy. By the time the survivors have hatched, fed, grown, migrated downstream, and transformed into smolts, most of the initial 3,000 offspring have died and left 20,800 kJ of assimilated energy in the ecosystem: 2,916 carcasses containing 6,869 kJ and 13,931 of respired heat energy.
2. juveniles in the ocean: The 84 surviving smolts have assimilated 7,423 kJ of food, expended 5,239 kJ on respiration, and amassed 2,184 kJ of body energy that they take with them to sea. There they feed and swim almost continuously, assimilating 505,508 kJ and expending 343,286 kJ on respiration. An additional 80 juveniles die in the ocean, leaving 388,016 kJ of assimilated energy in the ecosystem: 162,226 of biomass in their carcasses and 225,790 kJ of respired heat energy.
3. migration and breeding: After two years at sea, body energy has increased about 30 times. The four survivors contain 70,506 kJ of biomass energy when they enter fresh water to start the return migration. Essentially all of this energy is dissipated in fresh water: 54,408 as respiration, 11,280 kJ in the four carcasses, 3,880 kJ in the new clutch of 3,000 eggs, and 938 kJ in sperm.
4. production and assimilation: We estimate the production and assimilation of the cohort, *P*_*cohort*_ and *A*_*cohort*_, of all offspring of an average female up until a given stage, *S*_*ref*_, as

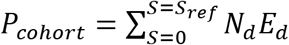

and

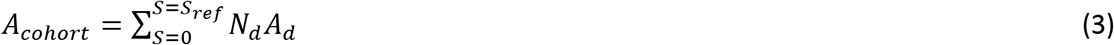

and where *N*_*d*_ is the number of offspring dying before stage, *S*, and *E*_*d*_ and *N*_*d*_ are respectively, total body energy content (production) and cumulative assimilation (production plus respiration) of those offspring when they died. Production efficiency of juveniles is high (32% of assimilation in fresh water and 46% in the ocean), but at the end of the life cycle only 5% the total energy assimilated by the cohort over one generation remains in fresh water.

**Table 2.** Impact of a cohort of Babine Lake sockeye salmon on fresh water and marine ecosystems. This accounting uses the MLT (Table 1) to quantify the input of food energy from assimilation (green) and its allocation to biomass production: growth and gametes (blue) and respiration (red).

| Ecosystem | Fresh water |  |  | Marine |  |  | Fresh Water |  |
| --- | --- | --- | --- | --- | --- | --- | --- | --- |
|  | Dead | Alive | Total | Dead | Alive | Total | Total | Net flow |
| Assimilation | 20800 | 7423 | 28223 | 88016 | 188002 | 576018 | 0 | 604241 |
| Respiration | 13931 | 5239 | 19170 | 225790 | 117496 | 343286 | 41525 | 403980 |
| Production | 6869 | 2184 | 9053 | 162227 | 70506 | 232733 | 0 | 200261 |
| Imported biomass |  |  |  |  |  | 2184 | 70506 |  |
| 4 Carcasses |  |  |  |  |  |  | 24162 |  |
| Clutch |  |  |  |  |  |  | 3880 |  |
| Sperm |  |  |  |  |  |  | 938 |  |
| Biomass left in fresh water |  |  |  |  |  |  | 28980 |  |
| Production efficiency | 0.23 | 0.21 | 0.22 | 0.29 | 0.26 | 0.28 | 0.0 | 0.23 |

#### Contribution of sockeye salmon to ecosystem energetics

Table 2 uses the energy budget of the cohort (Fig. 4B) and the MLT (Table 1) to quantify the impact of salmon on freshwater and marine ecosystems. Of the food energy assimilated by juveniles in fresh water, 92% is dissipated as heat of respiration and biomass of carcasses and 8% is transported into the ocean by the newly transformed smolts. Of the energy assimilated by juveniles in the ocean, 88% is dissipated in the marine ecosystem as respiration and mortality and 12% is transported into streams and lakes in the bodies of the four returning adults. The majority of this stored body energy (59%) is expended on respiration. The biomass in the carcasses, eggs and sperm is essentially all consumed by predators and decomposers; it amounts to only 41% of the energy brought in from the ocean and only 5% of the total energy assimilated by the cohort over one generation.

The production efficiency of a cohort or species population, *T*_*cohort*_, which Brett (1986) called the “growth efficiency”, is

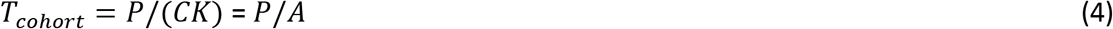

where *P* is the rate of biomass production, *C* is the rate of biomass consumption, and *K* is the energy conversion efficiency for a consumer (e.g., salmon as predator). So *T*_*cohort*_ indexes the efficiency of converting assimilated food eaten by salmon into salmon biomass that is ultimately consumed by other organisms – predators, scavengers and decomposers – in the freshwater and marine ecosystems. We calculated *T*_*cohort*_ (Table 1) using Brett’s (1986) estimate that *K* = 0.30 and assuming 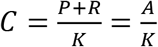. The trophic efficiency of juveniles is somewhat lower in fresh water (0.22) than in the ocean (0.28). After going without feeding after re-entering fresh water, the overall trophic efficiency over the entire life cycle is 0.23. For Babine Lake sockeye population, this efficiency is 23%.

### Discussion: broader implications

#### Physiology, ecology and evolution of salmon

*Oncorhynchus nerka* is a an exemplar of how natural selection has created a unique suite of physiological, behavioral and ecological traits (Quinn, 2018). These special features beg for explanation in terms of phylogenetic history and natural selection playing out in temporally and spatially varying environments. They are typically addressed by different disciplines and presented in specialized publications. The MLT provides a more synthetic explanation from the perspective of energetics. By combining the demography of a traditional life table with the physiological ecology of a balanced energy budget, the MLT applies first principles of physics and biology to integrate and synthesize many aspects of population biology, physiological ecology, and ecosystem energetics. These topics are typically addressed by different disciplines and presented in specialized publications. The MLT for Babine Lake sockeye emphasizes how traits at different levels of organization contribute to a unified big-picture perspective on salmon biology.

#### Energetics of migration

Several features of the MLT reflect the central role of migration. Very little energy is assimilated and produced by the fresh water stages, even though they account for most of the individuals and about half of the four-year life cycle. Despite the large clutch of eggs, because of small size and high mortality of the juvenile freshwater stages, they account for only a small fraction of cohort assimilation and biomass. Nearly all of the biomass energy produced over the life cycle (98% for a surviving breeder and 96% for an entire cohort) is acquired during feeding and growth in the ocean. The last stages of the life cycle are fueled by stored body energy, transported into fresh water and expended on the respiration for migration and breeding. By the time they spawn, two returning adults have expended 74% of their body energy, leaving only 19% in depleted carcasses and 7% in gametes.

#### Sockeye and kokanee

Comparisons between closely related salmon populations provide additional insights. As indicated by fossil history and shared adaptations for breeding in fresh water, the ancestral salmonids lived in streams and lakes (Hoar, 1976). Multiple lineages subsequently independently evolved life histories in which juveniles migrate to sea, grow to maturity, and migrate back to fresh water to breed (McCormick & Saunders, 1987). A consequence is a diversity of salmonid lifestyles, with closely related migratory and sedentary, anadromous and “land-locked” populations.

*Oncorhynchus nerka* is a good example. Sea run sockeye are widely distributed, with marine stages in the North Pacific and freshwater stages in surrounding lakes and streams. Another morpho-ecotype, known as kokanee, spends its entire life cycle in fresh water, with native populations in lakes scattered throughout the range and introduced populations more broadly distributed (Quinn, 2018). Kokanee and sockeye represent alternative stable evolutionary strategies: different combinations of traits that confer equal fitness. Sexually mature landlocked kokanee of the same age are smaller and less fecund (~700 g and ~800 eggs) than sea-run sockeye (~2,700 g and ~3,000 eggs) (USDA Forest Service, Lake Tahoe Basin Management Unit, Tahoe Heritage Foundation, 2015). By migrating to sea and feeding on the abundant plankton, anadromous populations were able to grow larger, produce more offspring, and occupy a distinctively different ecological niche. Despite such initial advantages that selected for the derived ocean-run ecomorphotype, however, kokanee and sockeye are now equally fit, as evidenced by the fact in some locations the two populations coexist and breed together but rarely hybridize. Compilation of a MLT for a population of kokanee would show quantitatively how the life history and energetics evolved in successful adaptation to its landlocked existence and reveal the tradeoffs resulting in equal fitness between kokanee and anadromous salmon. MLTs could also be used to predict effects of climate change, food restriction and other human activities on individual performance, cohort energetics, and ecosystem impacts of salmon and other ectotherms (see Huey & Kingsolver, 2019)

#### Ecosystem energetics

One consequence of the anadromous lifestyle is that migrating salmon transport large quantities of energy and materials from marine to freshwater and surrounding terrestrial ecosystems (Burger et al., 2012; Quinn, Helfield, Austin, Hovel, & Bunn, 2018; Schindler, Leavitt, Brock, Johnson, & Quay, 2005). Inorganic compounds in the bodies of salmon are major sources of nutrients for freshwater algae and land plants. A little more than half of the organic compounds (lipids, carbohydrates and proteins) are catabolized in aerobic respiration, generating ATP to fuel migration and reproduction, and releasing carbon dioxide, water and heat into the environment. A smaller but substantial quantity of the biomass is consumed by predators, scavengers and decomposers in freshwater and surrounding terrestrial ecosystems.

The ecological impact of these inputs is substantial. Each returning migrant transports only a small quantity of biomass, but the large body size and sheer numbers of migrants results in a significant net effect – a “marine subsidy” of energy and nutrients to freshwater and terrestrial ecosystems. When Brett did his studies in the late 1970s and early 1980s, the Skeena River sockeye was in approximate steady state and the annual runs of returning adults averaged about 3 million (DFO, 2003; Larkin & McDonald, 1968). Multiplying this by the 17,626 kJ of body energy per individual leaving the ocean gives about 53,000 megajoules of ocean production (or 1,500 tonnes of organic carbon) transported to freshwater and terrestrial ecosystems. For reference, this energy subsidy from a one-year run is equal to the annual net primary production, ~25 gC/m^2^, of approximately 6,000 ha of taiga forest (Gower et al., 2012).

In watersheds where the number of returning salmon have decreased due to human activities (see below), the reduced marine subsidies of energy and nutrients have caused substantial changes in freshwater and terrestrial ecosystems (Burger et al., 2012; Quinn et al., 2018; Schindler et al., 2005). There is also a “freshwater subsidy” to marine ecosystems due to juveniles (smolts) entering the ocean. Our MLT based on Brett’s data suggests that this is only about 3% of the marine subsidy (Naiman, Bilby, Schindler, & Helfield, 2002; Moore & Schindler, 2004; Scheuerell, Levin, Zabel, Williams, & Sanderson, 2005).

#### Use of wild salmon by humans

Few wild animals have been as important to humans as salmon. For thousands of years native fishers captured returning breeders in nets and traps at the mouths of rivers, preserved their flesh by drying and smoking, and budgeted the stored food to last through the lean months and years between good runs. Exploitation increased after the arrival of Europeans. Sockeye populations in the U.S. and Canada have declined precipitously due to overfishing and damming and pollution of rivers. Populations in Alaska and Asia are still abundant, due in part to supplementation from hatcheries and climate change (Ruggerone and Irvine 2018), although body sizes are declining (Oke et al., 2020). Ironically, one of the seriously depleted stocks is the Skeena River population that Brett studied. Annual returns fluctuated around 3 million until the late 1990s, then dipped to all-time lows. Now commercial, recreational and Native American fishing has been severely restricted (DFO, 2003; Hoekstra, 2017).

#### Theoretical implications

Recent advances in metabolic ecology raise the prospect for a general theory that uses energy as a fundamental currency to link tradeoffs in traits of individual organisms to their ecological and evolutionary manifestations – to show how anatomy, physiology and behavior affect energy budgets and in turn demography, population dynamics, community organization, and natural selection underlying biodiversity.

#### Comparison with dynamic energy budgets

MLTs are somewhat similar to the dynamic energy budgets of Kooijman and collaborators (DEBs: e.g. Kooijman & Kooijman, 2010; Sousa, Domingos, & Kooijman, 2008; Sousa, Domingos, Poggiale, & Kooijman, 2010). Both aim to contribute to a general theory of biological metabolism, using data to test theoretical predictions. Both have been applied to empirical model data on Pacific salmon (Nisbet, Jusup, Klanjscek, & Pecquerie, 2012; Pecquerie, Johnson, Kooijman, & Nisbet, 2011). The primary difference is that the DEB focuses on how underlying processes at molecular and cellular levels contribute to metabolic homeostasis at the individual organism level, whereas the MLT focuses on how metabolism at the whole-organism level affects demography of populations and energetics of ecosystems. Consequently, DEB models are usually have more details and require more parameter estimates, whereas MLTs incorporate just a few robust assumptions and parameters.

The two frameworks offer alternative, but not mutually exclusive theories applying biophysical laws to understand both species-specific and universal characteristics of living things. Biophysical laws that apply to all species limit how they acquire and allocate energy and constrain their life histories. The result is an interesting combination of universal and species-specific traits. On the one hand, all species are subject to: 1) the demographic constraint that in each generation at steady state parents are exactly replaced by offspring; 2) the energy balance constraint that all biomass energy assimilated from the environment is expended on survival, growth and reproduction; and 3) biological scaling laws that constrain for patterns of co-variation among traits. On the other hand, the diversity of living things offers examples of species with a wide variety of life histories: orders-of-magnitude variation in body size, growth and mortality, number and size of offspring, and reproductive mode (e.g., asexual and sexual, iteroparous and semelparous, degree of development at birth and subsequent parental care, etc.). Some of the universality and diversity predicted by existing metabolic theory is supported by empirical studies. Compilation of MLTs for species in diverse taxonomic and functional groups and comparisons among them offer exciting opportunities to test and extend metabolic theory. For example, an important component of energetic fitness is *F*, the fraction of cohort biomass production lost to mortality and consumed by other organisms in the ecosystem (Brown, Hall, & Sibly, 2018; Burger et al., 2019; Burger, Hou, Hall, & Brown, 2020; see below).

In principle, it is straightforward to construct a MLT for any species. Ideally, it would be put together by a single investigator or group of collaborators with first-hand experience working on diverse aspects of the biology. In practice, however, MLTs and the basic data necessary to compile are not available for more than a handful of well-studied species. The entries for a species in the increasingly available electronic databases suffer from the problems that often come from multiple studies of different populations and localities that define and measure the critical traits inconsistently. Our search of the literature for life tables and energy or mass balance diagrams generated many citations, but most are restricted to only part of the life cycle or lack critical data (e.g., many life tables do not give fecundity as a function of age).

Our best attempt at compiling MLTs based on body size for eight animals is in Burger et al., (2020). While this is a good start, it is important to note that most of these contain incomplete data and unverified assumptions. Comparison among species in this admittedly small sample suggests that *F*, the fraction of cohort biomass production reflects the tradeoff between number and size of offspring. Teleost fish that produce very large numbers of very small eggs have low values of *F* (e.g., 0.051 and 0.013 for salmon and Pollock, respectively), whereas mammals that supply parental care and rear a small number of relatively large offspring have much higher values (e.g., *F* = 0.35 and 0.25, for chimpanzee and brown bat, respectively) (Burger et al. 2020).

In conclusion, the MLT framework that Brett pioneered decades ago to synthesize information on life history, demography, energetics and physiology of sockeye salmon should be widely applicable to contemporary ecology. Most existing studies of metabolic ecology are based on extrapolating short-term measurements of respiratory metabolism – made most often in the laboratory (for basal, resting, active and maximal metabolic rates) and more rarely in nature (field metabolic rate). Brett compiled data from such short-term studies to put together a “life table of energetics” of sockeye salmon that serves as the foundation for the present paper. The MLT presented here quantifies energy gain from food and loss to heat and mortality, and energy allocation between respiration to generate ATP to fuel the costs of living and production of biomass energy passed on to the next generation in the form of offspring growth and parental gametes. So, it offers a synthetic picture of the central role of energy in the physiology, behavior, ecology and evolution of this one biologically distinct and economically important species. It provides an encouraging example of how the compilation of field data for more species in a MLT framework can allow us to evaluate and extend empirical and theoretical approaches to life history, ecology and evolution based in energetics (Brown et al., 2018; Burger et al., 2019; Burger et al. 2020). Doing so will has additional applications for sustainable resource management, conservation and predicting global change impacts on biodiversity.

## Acknowledgments

JRB acknowledges support from the Bridging Biodiversity and Conservation Science program in the Arizona Institutes for Resilience at the University of Arizona.

## Author contributions

All authors conceived the ideas and designed methodology; collected and analyzed the data; and wrote the manuscript including contributed critically to the drafts and gave final approval for publication.

## Notes

### Competing Interest Statement

The authors have declared no competing interest.

### Summary of Updates

Major revisions, new analyses and corrected errors in original data for egg size.

